# Rescue of two trafficking-defective variants of the neuronal glycine transporter GlyT2 associated to hyperekplexia

**DOI:** 10.1101/2021.03.24.436638

**Authors:** Andrés de la Rocha-Muñoz, Elena Melgarejo, Carmen Aragón, Beatriz López-Corcuera

**Affiliations:** Departamento de Biología Molecular Universidad Autónoma de Madrid. Spain; Centro de Biología Molecular “Severo Ochoa” Consejo Superior de Investigaciones Científicas-Universidad Autónoma de Madrid. Spain; IdiPAZ-Hospital Universitario La Paz, Universidad Autónoma de Madrid, Spain

**Keywords:** transport, glycine, hyperekplexia, mutation, chemical chaperone, calnexin

## Abstract

Hyperekplexia is a rare sensorimotor syndrome characterized by pathological startle reflex in response to unexpected trivial stimuli for which there is no specific treatment. Neonates suffer from hypertonia and are at high risk of sudden death due to apnea episodes. Mutations in the human *SLC6A5* gene encoding the neuronal glycine transporter GlyT2 may disrupt the inhibitory glycinergic neurotransmission and cause a presynaptic form of the disease. The phenotype of missense mutations giving rise to protein misfolding but maintaining residual activity could be rescued by facilitating folding or intracellular trafficking. In this report, we characterized the trafficking properties of two mutants associated with hyperekplexia (A277T and Y707C, rat numbering). Transporter molecules were partially retained in the endoplasmic reticulum showing increased interaction with the endoplasmic reticulum chaperone calnexin. One transporter variant had export difficulties and increased ubiquitination levels, suggestive of enhanced endoplasmic reticulum-associated degradation. However, the two mutant transporters were amenable to correction by calnexin overexpression. Within the search for compounds capable of rescuing mutant phenotypes, we found that the arachidonic acid derivative N-arachidonoyl glycine can rescue the trafficking defects of the two variants in heterologous cells and rat brain cortical neurons. N-arachidonoyl glycine improves the endoplasmic reticulum output by reducing the interaction transporter/calnexin, increasing membrane expression and improving transport activity in a comparable way as the well-established chemical chaperone 4-phenyl-butyrate. This work identifies N-arachidonoyl glycine as a promising compound with potential for hyperekplexia therapy.

## 1. Introduction

Hyperekplexia or startle disease (OMIM 149400) is a rare sensorimotor syndrome characterized by pathological startle reflex of lower threshold, higher intensity, and resistance to habituation (Brown et al. 1991). Patients present neonatal hypertonia that may cause apnea episodes, which puts them at risk of brain damage or even sudden infant death. The adults suffer very disabling motor alterations and recurrent unprotected falls throughout their entire life (Andermann et al. 1980). The disease has no causal treatment but symptomatic pharmacotherapy with benzodiazepines such as clonazepam, which may produce side effects, sometimes very limiting, such as sedation (Dreissen and Tijssen 2012).

Hyperekplexia is caused by disruption of the inhibitory glycinergic neurotransmission, which coordinates the startle reflex, among other functions (Bakker et al. 2006). The main cause of the disease are mutations in the strychnine-sensitive glycine receptor, a glycine-gated chloride channel that responds to neurotransmitter released from glycinergic interneurons. Glycine receptor activation hyperpolarizes the postsynaptic membrane suppressing excitatory postsynaptic potentials and causing inhibition (Legendre 2001; Langlhofer et al. 2020). The second most common cause of hyperekplexia are mutations in the *SLC6A5* gene, encoding the presynaptic glycine transporter GlyT2 that cotransports glycine to the presynaptic terminal together with Na^+^-and Cl^−^ (Thomas et al. 2013). The neuronal GlyT2 is a crucial protein for synaptic glycine recycling in inhibitory glycinergic synapses, a necessary step for synaptic vesicle refilling (Rousseau, Aubrey, and Supplisson 2008). Transporter loss of function abolishes glycinergic neurotransmission (Gomeza et al. 2003). GlyT2 mutations associated to hyperekplexia include alterations of residues essential for the transport activity, changes in amino acids involved in conformational changes needed for the translocation cycle and also mutations preventing transporter trafficking to the plasma membrane (Kitzenmaier et al. 2019; Dafsari et al. 2019). In this latter group, there are GlyT2 mutants that maintain residual or even significant activity for which facilitation of the intracellular trafficking could rescue the phenotype.

GlyT2 is a 12 transmembrane domain (TM) protein belonging to the SLC6 family of neurotransmitter transporters, which also includes the transporters for GABA and monoamines (Rudnick et al. 2014). The structural hallmark of this family includes two topologically inverted repeats of 5 TMs each that arrange forming two bundles: a scaffold bundle and a core bundle. During the translocation cycle, the core bundle rocks against the scaffold bundle making accessible the central substrate site to one or another side of the membrane (Jardetzky 1966). In this way, the protein goes through at least three conformational states: outward-open, occluded and inward-open, all of them crystallized (Drew and Boudker 2016; Shahsavar et al. 2021).

GlyT2 is synthetized in ribosomes associated to the endoplasmic reticulum (ER) and co-translationally translocated to the ER membrane so that intracellular N- and C-termini lay in the cytoplasm while protein folding and modification takes place assisted by ER chaperones, such as calnexin (CNX) (Arribas-Gonzalez et al. 2013). Properly folded proteins are transported from the ER to the Golgi apparatus in vesicles coated by coatomer protein II (COPII) by using the Sec24D adaptor or, if permanently unfolded, they are exported for ER-associated degradation (ERAD) (Needham, Guerriero, and Brodsky 2019). The biogenesis of plasma membrane proteins can be facilitated by using chemical chaperones such as 4-phenylbutyric acid (PBA), which can rescue wild-type GlyT2 from a dominant negative trafficking mutant associated to hyperekplexia (Arribas-Gonzalez et al. 2015). PBA seems to interact with exposed hydrophobic patches in the proteins (Cortez and Sim 2014). In addition, more specific pharmacochaperones that interact with specific folding intermediaries or loosen the ER quality control may also have a folding promoting action (Bhat, Newman, and Freissmuth 2019).

In this report, we focus on the trafficking properties of two missense hyperekplexia mutations (A275T and Y705C) that had been previously shown to impair transporter function (Carta et al. 2012; Gimenez et al. 2012). The substitution in A275T mutant affects crucial residues for sodium binding and is detrimental for glycine transport, although protein trafficking was not reported to be altered (Carta et al. 2012). Mutant Y705C, one of the scarce mutations with dominant inheritance, introduces a cysteine that alters proton dependence of transport, besides affecting transporter maturation and function. Though, the intracellular compartment were the transporter was arrested remains elusive (Gimenez et al. 2012). In this report, we have identified the trafficking defects of the rat versions of A275T and Y705C mutants (A277T and Y707C, rat numbering). Besides, through searching for compounds capable of rescuing their phenotype, we found that the arachidonic acid derivative N-arachidonoyl glycine (NAG) can rescue the functional defects of the two variants in rat brain cortical neurons.

## 2. Material and methods

### 2.1. Materials

Female Wistar rats were bred under standard conditions at the Centro de Biología Molecular Severo Ochoa (CBMSO) in accordance with procedures approved in the Directive 2010/63/EU of the European Union with approval of the Research Ethics Committee of the Universidad Autónoma de Madrid (Comité de Ética de la Investigación UAM, CEI-UAM). Rabbit and rat antibodies against N-terminus of GlyT2 were generated in house (Zafra et al. 1995). Other primary antibodies used were: anti-transferrin receptor (Invitrogen, #13-6800), anti-E cadherin (a gift from Amparo Cano, UAM), rabbit anti-calnexin (StressMarq Biosciences, SPC-108), mouse anti-calnexin (BD Transduction Laboratories, clone 37), anti-ubiquitin (Santa Cruz, sc-8017, clone P4D1), anti-α-tubulin (Sigma-Aldrich, clone T-6074), anti-Myc (Cell Signaling, #2276) and anti-transferrin receptor (Invitrogen). For the screening of compounds with the potential to rescue GlyT2 defective phenotypes the following chemicals were used: bupropion hydrochloride (Sigma Aldrich), ibogaine hydrochloride (LGC Standards), ALX1393 (Santa Cruz Biotechnology), N-arachidonoyl glycine (Cayman chemicals) and 4-phenylbutirate (Sigma Aldrich). All other chemicals used were from Sigma Aldrich unless otherwise noticed. Neurobasal medium and B27 supplement were purchased from Invitrogen.

### 2.2. Plasmid constructs

The myc-Sec24D construct in pcDNA3 (Invitrogen) was generously provided by Francisco Zafra (Centro de Biología Molecular Severo Ochoa. Madrid, Spain). GlyT2 was subcloned into pcDNA3 and the GlyT2 mutants were constructed by site-directed mutagenesis using the QuikChange kit (Stratagene). Plasmids from two independent *Escherichia coli* colonies were transfected into eukaryotic cells and [^3^H]glycine transport was measured in the cells for verification. The CNX cDNA clone (IMAGE number 2582119) in pCMV.SPORT6 was purchased from Source Bioscience Lifesciences (Arribas-Gonzalez et al. 2013).

### 2.3. Cell lines and protein expression

COS7 cells (American Type Culture Collection) were grown in Dulbecco’s modified Eagle’s medium (DMEM) supplemented with 10% fetal bovine serum (FBS). MDCK II (American Type Culture Collection) cells were grown in minimum essential medium (MEM) supplemented with 10% FBS. Cells were maintained in incubators at 37 °C and 5% CO_2_. Turbofect Transfection Reagent or Lipofectamine 2000 (ThermoFisher Scientific) were used to transfect COS7 cells and MDCK II cells, respectively. Transfections were performed following manufacturer’s protocol, adding the reagents and DNA to the culture media in the absence of supplements (2 μl of reagent per μg of DNA). Then, cells were maintained for 48 h at 37 °C until use.

### 2.4. Primary cultures of cerebral cortex

Primary cultures of embryonic cortical neurons were prepared as described previously (Arribas-Gonzalez et al. 2015). Briefly, the cortex of Wistar rat fetuses was obtained on the 18^th^ day of gestation (E18) and the tissue was mechanically disaggregated in Hanks’ balanced salt solution (HBSS; Invitrogen) containing 0.25% trypsin (Invitrogen) and 4 mg/ml DNase (Sigma). Cells were plated at a density of 500,000 cells/well in 6 well plates (Falcon) and incubated for 4 h in DMEM + 10% FCS, containing: glucose, 10 mM; sodium pyruvate, 10 mM; glutamine, 0.5 mM; gentamicin, 0.05 mg/mL; streptomycin, 0.1 mg/mL; and penicillin G, 6×10^−5^ mg/mL. After 4 hours the buffer was replaced with Neurobasal/B27 culture medium containing glutamine (0.5 mM, 50:1 by volume; Invitrogen) and 3 days later cytosine arabinoside (2.5-5 × 10^−3^ mM) was added to inhibit further glial growth. For transfection, neurons that had been maintained in vitro for 5 to 7 days (DIV 5 to 7), were incubated with 2 μg of total DNA mixed with 4 μl of Lipofectamine 2000 reagent (Invitrogen) following supplier instructions. GlyT2 glycine transport and membrane expression were measured after 48h in culture.

### 2.5. [^3^H]glycine transport assays

GlyT2 transport activity was measured as described previously (Benito-Munoz et al. 2018). Briefly, cells were washed and incubated at 37 °C in phosphate buffered saline (PBS) containing 2 μCi/ml [^3^H]-glycine (1.6 TBq/mmol; PerkinElmer Life Sciences) isotopically diluted to reach 10 μM final glycine concentration. The reactions were terminated by aspiration after 10 min, followed by PBS wash. To determine GlyT2-specific glycine transport, glycine accumulation by non-transfected cells (mock) was also measured and subtracted from that of GlyT2-expressing cells. In primary neurons, the transport medium contained the GlyT1 antagonist NFPS (10 μM), and the basal uptake was measured in the presence of the GlyT2-selective inhibitor 1 μM ALX1393 (de la Rocha-Muñoz et al. 2019). Transport activity was normalized to the protein concentration and expressed in pmol of Gly/μg of protein/ 10 min.

### 2.6. Electrophoresis and Western blotting

Protein samples were separated by SDS-PAGE using a 4% stacking gel and 6 or 7.5% resolving gels. The samples were transferred to nitrocellulose with a semi-dry transfer system (Life Technologies Inc.: 65V/gel, 90 min). Membranes were blocked for 30 min with 5% milk in PBS containing 0.1% Tween 20 at 25 °C. The membranes were probed overnight at 4 °C with the desired primary antibody: anti-GlyT2 (rabbit 1:1,000); anti-GlyT2 (rat, 1:500); anti-Myc (mouse, 1:1,000) or anti-ubiquitin (mouse, 1:200). After several washes, the antibodies bound were detected with peroxidase coupled anti-rat (1:8,000; Sigma), anti-rabbit (1:8,000; Bethyl) or anti-mouse IgG (1:8,000; ThermoFisher Scientific) which were visualized by enhanced chemiluminescence (ECL; Amersham Corp.). Subsequently, the antibodies were stripped from the membrane (Thermo Scientific), which was re-probed with anti-tubulin (mouse, 1:2,000), anti-CNX (mouse, 1:1,000); anti-transferrin receptor (mouse, 1:500) or anti-GlyT2 (rat 1:500) as loading controls. Antibody binding was detected with a peroxidase-coupled anti-mouse or anti-rat IgG. Protein bands visualized using film exposures in the linear range were imaged using a GS-900 calibrated imaging densitometer (Bio-Rad) and quantified using Image Lab Software (Bio-Rad).

### 2.7. Surface biotinylation

As described (Arribas-Gonzalez et al. 2015). Cells expressing wild-type or mutant GlyT2 grown in 6 well plates (Nunc) were washed and labeled with Sulfo-NHS-Biotin (1.0 mg/ml in PBS; Pierce) at 4 °C, a temperature that blocks membrane trafficking of proteins. After quenching with 100 mM L-lysine to inactivate the free biotin, protein concentration was measured and cells were lysed with radioimmunoprecipitation assay (RIPA) buffer. A portion of the lysate was saved as total protein fraction (T), and the remainder was incubated with streptavidin-agarose beads for 2h at room temperature (RT) with rotary shaking. After centrifugation, the supernatant was kept as non-biotinylated fraction (intracellular proteins). The agarose beads recovered were washed 3 times with 1 ml RIPA buffer and bound proteins (surface-biotinylated) were eluted with 2x Laemmli buffer (65 mM Tris, 10% glycerol, 2.3% SDS, 100 mM DTT, 0.01% bromophenol blue) for 10 min at 75 °C. Samples were resolved in SDS-PAGE and analyzed in Western blots using transferrin receptor (TfR) immunoreactivity as loading control.

### 2.8. Carbohydrate modification

COS7 cells expressing GlyT2 or the desired mutants were lysed in 1x lysis buffer (150 mM NaCl, 50 mM Tris-HCl [pH 7.4], 5 mM EDTA, 1% Triton-X100, 0.1% SDS, 0.25% deoxycholate sodium, 0.4 mM phenylmethylsulfonyl fluoride (PMSF) and 4 µM pepstatin) and digested with the chosen endoglycosidase (PNGase F, New England Biolabs; or Endoglycosidase H, Roche) in a small volume of the appropriate buffer, according to the manufacturer’s instructions. These cell extracts were then resolved by SDS-PAGE and analyzed in Western blots (Arribas-Gonzalez et al. 2015).

### 2.9. Coimmunoprecipitation assays

Performed as described (Arribas-Gonzalez et al. 2015). Transfected COS7 cells were washed twice with PBS, scrapped off the plates in 150 mM NaCl, 50 mM Tris-HCl [pH 7.4], 0.4 mM PMSF and 4 µM pepstatin and the desired amount of protein (Bradford method, Bio-Rad) was solubilized for 90 min at RT in in PBS containing 0.2% Nonidet P-40 substitute, 0.4 mM PMSF and protease inhibitor Sigma cocktail (coIP buffer). After 15 min centrifugation at 10,000 x g, an aliquot of the lysate was retained to measure the total protein content and the remainder was precleared by adding 20 μl of 50% protein G-agarose (PGA, Invitrogen) in coIP buffer for 30 min at 4 °C with rotation. After centrifugation, the supernatants were incubated overnight at 4 °C with 2 μg of the desired primary antibody (rabbit anti-CNX or rabbit anti-GlyT2). Controls with no antibody were also included. Subsequently, 20 μl of PGA beads in coIP buffer was added to the samples and after incubating for 90 min at RT, the beads were washed 3 times for 5 min with coIP buffer. The bound proteins were then dissociated from the beads by heating at 75 °C for 10 min in 2x Laemmli buffer, resolved by SDS-PAGE (7.5%) and analyzed in Western blots. CNX and GlyT2 immunoreactivities used as loading control were obtained by probing with mouse and rat antibodies respectively.

### 2.10. Ubiquitination assay

As indicated in (de la Rocha-Muñoz et al. 2019). COS7 cells were washed twice with PBS at 4 °C, harvested using Ub buffer: 50 mM Tris, 150 mM NaCl and 50 mM N-ethylmaleimide and Sigma cocktail of protease inhibitors (1:200), and cell protein content was determined (Bradford). Samples containing equal amounts of protein were centrifuged and cell pellets were resuspended in 90 microliters of Ub buffer. Then, sodium dodecyl sulphate (SDS) was added to a final concentration of 1% and samples were incubated for 10 min at 95 °C to disrupt protein interactions. Then, 4% Triton X-100 was added to reach 1% and samples were diluted by adding 1 ml of Ub buffer containing 1% Triton X-100. After 30 min of rotary shaking at 4 °C, lysates were precleared at 4 °C with 50% PGA in Ub buffer during 30 min. The supernatants were incubated overnight with anti-GlyT2 antibody at 4°C and then incubated with 50% PGA for 90 min at RT. PGA beads were washed 3 times with ice-cold Ub buffer and bound proteins eluted in 2x Laemmli buffer. Samples were resolved in 6% SDS-PAGE gels and subjected to western blot with the ubiquitin-specific antibody.

### 2.11. Immunofluorescence

Cells expressing wild-type or mutant transporters growing on coverslips were washed 3 times with PBS and fixed with methanol at −20 °C for 15 min (Jimenez et al. 2011). Cells were permeabilized in PBS containing 0.1% Triton X-100 at RT for 10 min and PBS washed. Then, cells were incubated with 2% FBS, 2% bovine serum albumin (BSA) and 0.2% bovine gelatin in PBS (blocking solution) for 1 h. After washing, cells were incubated with primary antibodies (rabbit anti-GlyT2 and rat anti-E-cadherin or rat anti-GlyT2 and rabbit anti-CNX, 1:200 each) diluted in PBS containing 10% blocking solution for 90 min at RT. Cells were washed and incubated with the secondary antibodies during 60 min at RT. After a final wash, cells were mounted on microscope slides using Mowiol and dried overnight in the darkness at RT. Samples were stored at 4 °C before visualization in an inverted confocal microscope LSM 510 (Zeiss). Colocalization degree of different probes was quantified with ImageJ, using Mander’s overlap coefficient included in JACoP plugin (Manders, Verbeek, and Aten 1993).

### 2.12. Data analysis and statistics

All data and statistical analyses were performed using GraphPad Prism (GraphPad Software). Kruskal-Wallis test was used to compare multiple groups, with subsequent Dunn’s post-hoc test to determine the significant differences between samples. Kolmogorov-Smirnov and Mann-Whitney U tests were used to compare two separate groups. p values are denoted through the text as follows: *p < 0.05; **p < 0.01; ***p < 0.001; ****p < 0.0001; p < 0.05 or lower values were considered significantly different. Thorough the text, the Box whisker plots represent median (line), mean (+), 25–75 percentile (box), 10–90 percentile (whisker), 1–99 percentile (X) and min - max (−) ranges.

## 3. Results

### 3.1. Two GlyT2 variants with missense mutations exhibit decreased surface expression and ER retention

Mutant A277T has a point substitution in TM3 within a region very close to the Na3 site (Benito-Munoz et al. 2018). When the transporter was expressed in COS7 cells, the glycine transport activity was reduced to 50.2 ± 2.3%, as compared to the wild-type (Fig.1A, n = 79, **** p <0.0001). This could be attributed to altered dependence on Na^+^ and glycine (Supplementary Fig. S1), in agreement to previous reports for the human A275T (Carta et al. 2012). On the other hand, mutant Y707C contains a substitution in TM11 and showed reduced transport activity (55.3 ± 1.4%, Fig.1A, n = 72, **** p <0.0001), when expressed in COS7. It has been proven that the introduced cysteine disrupts the structural disulphide bond present in the external side of the transporter protein hindering its maturation along the secretory pathway and reducing membrane expression and function. However, transporter trafficking was not (Carta et al. 2012) or incompletely studied in these mutants (Gimenez et al. 2012).

**Figure 1.**
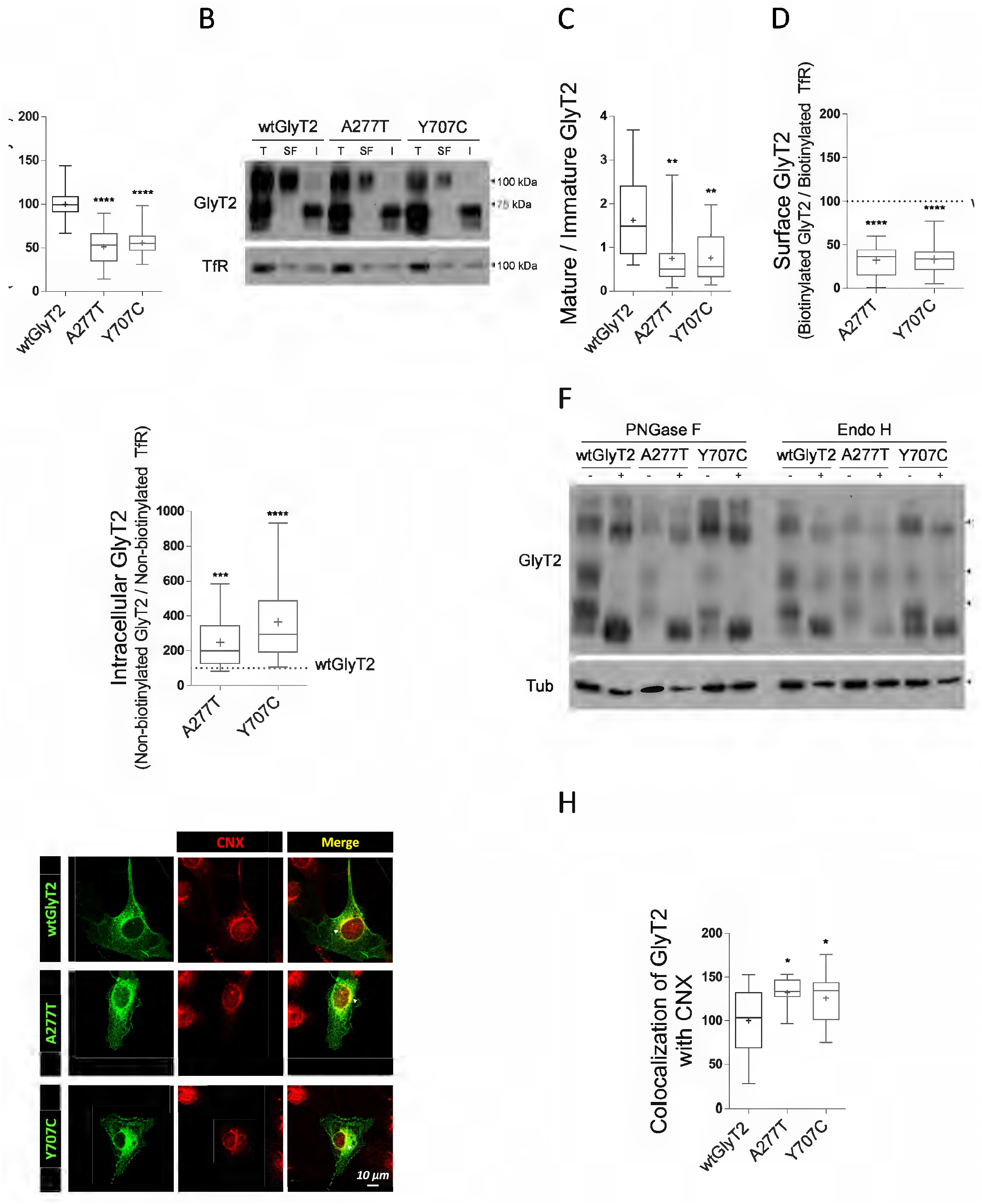
A277T and Y707C mutants have reduced uptake activity caused by an altered trafficking. **A)** COS7 cells transiently expressing wtGlyT2 (wild-type), A277T or Y707C were subjected to [^3^H]-glycine transport assays. Glycine transport shown is normalized against wtGlyT2. ****p < 0.0001, using Dunn’s multiple comparison test. n (wtGlyT2) = 88, n (A277T) = 79, n (Y707C) = 72. **B)** COS7 cells transiently transfected as in (A) were subjected to surface biotinylation: surface biotinylated (SF), intracellular-not biotinylated (I) and total (T) transporter fractions from the lysate were analyzed by western blot. Transferrin receptor (TfR) immunoreactivity was used as loading control. **C)** Intensity ratio between mature (100 kDa) and immature bands (75 kDa) of the transporter total fraction. **p (A277T) = 0.0028, **p (Y707C) = 0.0027; using Dunn’s multiple comparison test. n (wtGlyT2) = 19, n (A277T) = 14, n (Y707C) = 16. **D)** Quantification of transporter biotinlytated fraction using TfR as loading control (percentage to the corrected signal of wtGlyT2, indicated by the dashed line). ****p < 0.0001; using Dunn’s multiple comparison test. n (wtGlyT2) = 15, n (A277T) = 15, n (Y707C) = 14. **E)** Quantification of transporter intracellular fraction using TfR as loading control (percentage to the corrected signal of wtGlyT2, indicated by the dashed line). ***p (A277T) = 0.0001, ****p (Y707C) < 0.0001; using Dunn’s multiple comparison test. n = 4. **F)** COS7 cells transiently transfected as in (A) were lysed and treated overnight with the vehicle alone (endoglycosidase buffer, -) or with the indicated endoglycosidase (+) in denaturing conditions and then resolved by SDS-PAGE. **G)** COS7 cells transiently transfected as in (A) were immunolabeled for GlyT2 (green) and the endoplasmic reticulum marker CNX (red). Arrowheads (△) indicates areas showing most colocalization. **F)** Colocalization of the transporters with CNX was quantified using Mander’s overlap coefficient and normalized against wtGlyT2. *p (A277T) = 0.0381, *p (Y707C) = 0.0384; using Holm-Sidak’s multiple comparison test. n (wtGlyT2) = 19, n (A277T) = 8, n (Y707C) = 16.

In order to study the trafficking properties of the two mutants, we subjected transporter-expressing COS7 cells to surface biotinylation. In these experiments, surface proteins were biotin-labelled and pelleted by binding to streptavidin beads. Aliquots of total proteins, beads-bound (surface) proteins and unbound (intracellular) proteins were subjected to western blot (WB) for detection of GlyT2. In the total fraction, monomeric GlyT2 appears as a doublet of two bands. One band, about 75 kDa, corresponds to the immature carrier residing in ER and presents incomplete glycosylation. The 100 kDa band is the fully glycosylated mature form (Fig. 1B)(Arribas-Gonzalez et al. 2013). Since the two transporter forms represent different maturation states of GlyT2 along the secretory pathway, the ratio mature (100 kDa) to immature (75 kDa) protein is informative on possible alterations in transporter biogenesis. For the two variants, this ratio was around 55% with respect to the wild-type (Fig. 1C, n (A277T) = 14, ** p = 0.0028; n (Y707C) = 16, ** p = 0.0027), indicating an altered maturation along the secretory pathway. On the other hand, in the surface-biotinylated fractions only the 100 kDa band was detected, as corresponds to the plasma membrane-residing GlyT2, able to perform the transport of glycine (Fig. 1B). The amount of transporter in the surface-biotinylated fraction was clearly decreased for the mutants as compared to the wild-type, being 34.6 ± 4.7%, n (A277T) = 15 and 33.6 ± 5.0%, n (Y707C) = 14 (**** p <0.0001)(Fig. 1D). Finally, the intracellular-unbound fraction was enriched in the immature form and specially in the mutants (Fig. 1E). A277T reached 248.0 ± 36.7% (n = 4, *** p = 0.0001) with respect to wild-type, whereas Y707C reached 365.8 ± 59.7% (n = 4, **** p < 0.0001). These results indicate the mutants have an intracellular retention problem that hinders delivery to the plasma membrane. These data indicate for the first time A277T mutation alters transporter trafficking along the secretory pathway and, on the other hand, are in agreement with previous reports on Y705C traffic (Gimenez et al. 2012).

Since GlyT2 mutants seemed to have defects in their maturation, we next studied the glycosylation state of both variants to determine if there were differences in their location in the secretory pathway (Fig. 1F). As previously shown (Arribas-González et al. 2015), peptide: N-glycosidase F (PNGase F) completely removed the N-linked glycans from both 75 and 100 kDa GlyT2 forms, yielding a 60 kDa band that corresponds to the non-glycosylated protein core. Besides, endoglycosidase H (Endo H) only affected the immature 75 kDa form and not the 100 kDa protein. The 75 kDa protein, located in ER, is sensitive to Endo H since it contains high mannose oligosaccharides. However, the 100 kDa form is Endo H-resistant, as it has exited the ER and undergone oligosaccharide processing in the Golgi. As Figure 1F shows, there were not differences in the sensitivity of the 75 and 100 kDa forms to PNGase F and Endo H for both variants as compared to wild-type, indicating that A277T and Y707C mutants are located in the same compartments than wild-type and undergo similar processing along the secretory pathway. Nevertheless, mutant Y707C, shows a double band in the untreated lane indicating Y707C has a glycosylated protein fraction (upper band) and a non-glycosylated fraction (lower band). The lower/non-glycoslylated band is more prominent in Y707C than in the wild-type lane (this is better observed in the Endo H blot than in the PNGase F blot), indicating that this mutant is retained in the ER.

In order to confirm ER retention by the mutants, we measured the degree of transporter colocalization with the ER chaperone calnexin (CNX), usually used as ER marker (Fig. 1G, H). For these experiments, the transporters were expressed in COS7 cells, and double immunofluorescence against GlyT2 and endogenous CNX was performed. Quantification using Manders overlap coefficient (Manders, Verbeek, and Aten 1993) indicated mutant distribution overlapped with CNX about 30% more than the wild-type (Fig. 1H, n (A277T) = 8, *p = 0.0381 and n (Y707C) = 16, *p= 0.0384). This points to a higher mutant concentration in the ER as compared to the wild-type, confirming the variants associated with human hyperekplexia are retained in the ER.

To obtain complementary evidence about mutant defective trafficking, the two GlyT2 variants were expressed in MDCK II cells and subjected to surface-biotinylation as above (Supplementary Fig. S2). As Supplementary Fig. S2A shows, WB of MDCK II total fraction presents a GlyT2 band pattern similar to that observed in COS7 cell lysates, with two monomeric forms around 75 and 100 kDa corresponding to different maturation states of the carrier. As in COS7 cells, the surface expression of GlyT2 mutants was decreased about 50% compared to wild-type (Supplementary Fig. S2B). Additionally, plasma membrane expression of the mutants was studied by double immunofluorescence with GlyT2 antibodies and antibodies against the endogenous plasma membrane marker E-cadherin. The MDCK II cell line is very convenient for these assays because it visibly expresses E-cadherin when certain level of confluence is reached. We quantified the degree of colocalization of the mutants with E-cadherin using the Manders overlap coefficient (Manders, Verbeek, and Aten 1993), and found it was about 30% lower than that observed for the wild-type (Supplementary Fig. S2C, D). Specifically, the overlap degree was 69.8 ± 4.0% for A277T (n = 35, **** p <0.0001) and 68.1 ± 3.9%for Y707C (n= 32, **** p <0.0001). Besides, since typical GlyT2 apical sorting can be recapitulated in polarized MDCK II cells grown in transwells, we analysed whether GlyT2 mutants present also an asymmetric distribution. As Supplementary Fig. S2E shows, the two GlyT2 mutants were distributed along the apical domain of the MDCK II cells, although to a lesser extent than the wild-type transporter, which corroborates their increased intracellular retention already demonstrated in COS7 cells.

### 3.2. The two variants show increased interaction with calnexin and different ER export and ubiquitination capability

Several processes take place in the ER during the early stages of the trafficking along the secretory pathway. First, we studied the interaction of the mutants with CNX, whose role in GlyT2 quality control was previously described by our group (Arribas-Gonzalez et al. 2013). To do this, COS7 cells expressing the transporters were lysed under permissive conditions for protein-protein interactions, immunoprecipitated with an antibody against CNX and subjected to WB for GlyT2 detection. As we previously reported, CNX interacts with the immature fraction of GlyT2 (75 kDa), which is the one found in the ER (Arribas-Gonzalez et al. 2013). However, under the conditions used in this work, a nonspecific band overlapping with the immature fraction masked its appearance and prevented its quantification (Fig. 2A). Instead, we quantified the SDS-resistant high molecular weight forms (250 kDa) of the immature transporter, which we verified were quantitatively identical to the 75 kDa form and gave an accurate approximation to the proportion of immature transporter coimmunoprecipitated with CNX. As previously shown in Fig. 1F the 250 kDa band has endoglycosidase sensitivity identical to that of the 75 kDa form, being sensitive to PNGase F and Endo H, discarding the aggregate contains the mature 100 kDa GlyT2. Quantification revealed the two mutants coimmunoprecipitated with CNX in greater proportion than the wild-type transporter (Fig. 2A, B). The amount of A277T coimmunoprecipitated with CNX was almost 3 times higher than that of wild-type GlyT2 (289.1 ± 120.2%, n = 4, **p = 0.0091). Y707C coimmunoprecipitated with CNX about twice more than wild-type (188.5 ± 20.1%, n = 6, **p = 0.0084). The increased interaction with CNX of the mutants suggests the mutations promote folding defects and therefore ER retention.

**Figure 2.**
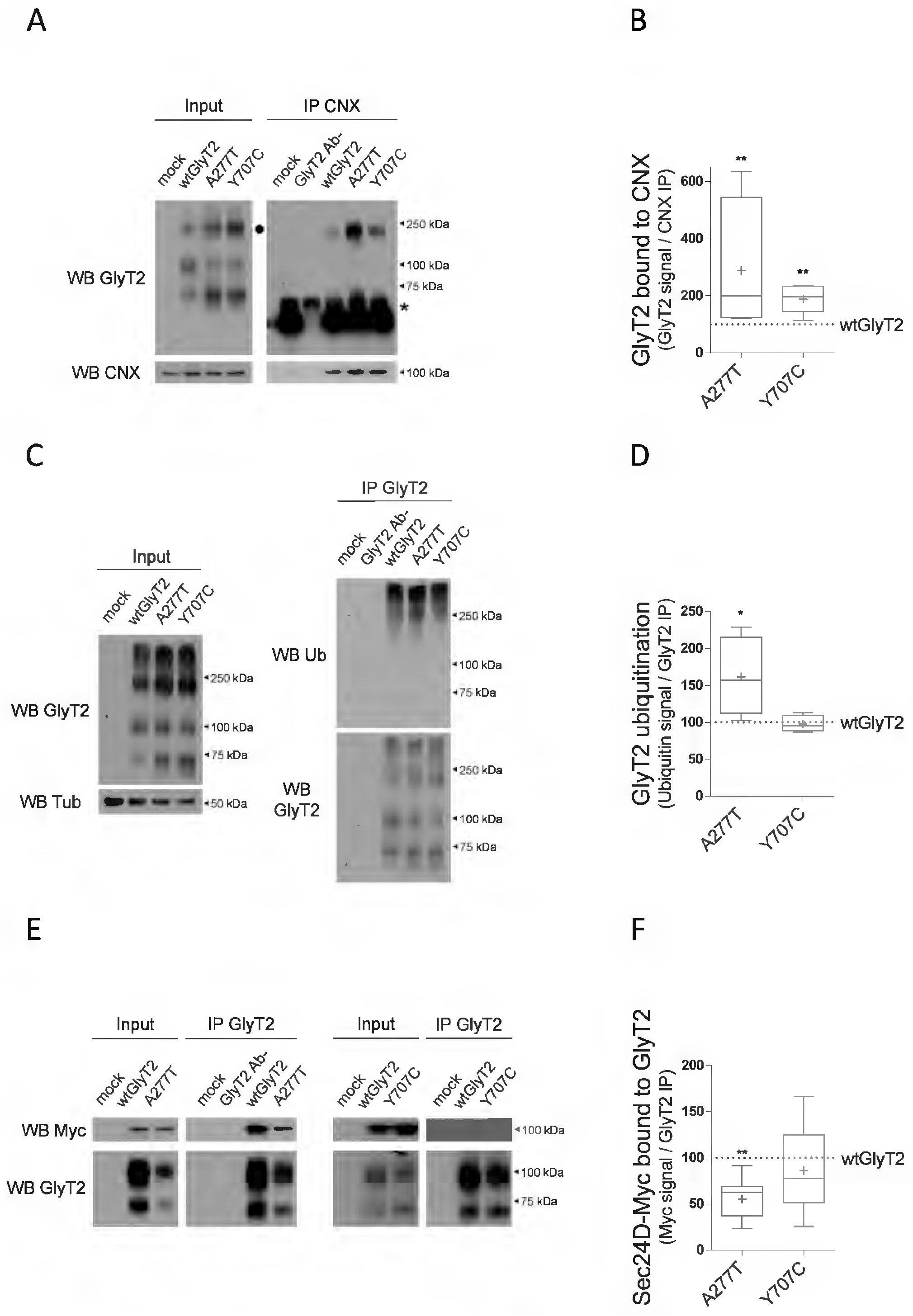
A277T mutation causes more severe retention of the transporter in the endoplasmic reticulum than Y707C mutation. **A)** COS7 cells lysates expressing wtGlyT2, A277T or Y707C were subjected to CNX immunoprecipitation using anti-CNX made in rabbit and the immunocomplexes were analyzed by western blot to detect CNX (using anti-CNX made in mouse) and GlyT2 (using anti-GlyT2 made in rat). 250 kDa aggregates of immature GlyT2 are indicated with a black circle (●); unspecific 50 kDa bands are indicated with an asterisk (*). **B)** Quantification of the amount of transporter bound to CNX normalized against the quantity of wtGlyT2 bound to CNX (indicated by the dashed line). **p (A277T) = 0.0091, **p (Y707C) = 0.0084; using Dunn’s multiple comparison test. n (wtGlyT2) = 6, n (A277T) = 4, n (Y707C) = 6. **C)** COS7 cells transiently transfected as in (A) were lysed, subjected to immunoprecipitation against GlyT2 and ubiquitination of the transporter was assayed by immunoblotting against ubiquitin (Ub). **D)** Quantification of transporter ubiquitination. Blots were probed against GlyT2 to normalize ubiquitination signal against the amount of GlyT2 immunoprecipitated in each case. Ubiquitination levels are normalized against ubiquitination levels of wtGlyT2 (indicated by the dashed line). *p (A277T) = 0.0401; using Dunn’s multiple comparison test. n = 4. **E)** COS7 cells transiently transfected as in (A) were lysed, subjected to immunoprecipitation against GlyT2 and the immunocomplexes were analyzed by western blot to detect Myc-Sec24D and GlyT2. **F)** Quantification of the amount of Myc-Sec24D bound to GlyT2 normalized against the amount of Myc-Sec24D bound to wtGlyT2 (indicated by the dashed line). **p (A277T) = 0.0019, ^ns^p (Y707C) = 0.1858; using Dunn’s multiple comparison test. n (wtGlyT2) = 7, n (A277T) = 7, n (Y707C) = 5. Mock: untransfected cell sample. GlyT2 Ab-: GlyT2 transfected but precipitation without antibody.

The permanence of the mutants in the ER and its enhanced interaction with CNX suggest there is a high proportion of misfolded transporters that must be degraded. Since misfolded proteins eliminated by ERAD are previously modified by ubiquitination (Hebert and Molinari 2007), we determined the level of ubiquitination of the mutants (Thibaudeau and Smith 2019). In order to visualize by WB the short half-life ubiquitinated proteins, we promoted their accumulation by inhibiting the proteasome with MG132 (Arribas-Gonzalez et al. 2013). Fig. 2C, D shows that the levels of ubiquitin associated with A277T are increased by about 60% (161.4 ± 26.7%; n = 4, * p = 0.0401), suggesting that the A277T mutant is targeted to ERAD to a higher extent than the wild-type transporter. In contrast, mutant Y707C showed wild-type levels of ubiquitination.

To monitor possible defects in ER export, we studied the interaction of the mutants with the Sec24D protein, an adaptor of the COPII complex that selects properly folded proteins for trafficking to the Golgi (Chiba, Freissmuth, and Stockner 2014). For this purpose, after co-expression of GlyT2 mutants with a Sec24D-Myc construct and immunoprecipitation with a GlyT2 antibody, the immunoreactivity against Myc was evaluated in GlyT2 immunocomplexes. The amount of Sec24D coinmunoprecipitated with A277T was 55.1 ± 8.7% of that bound to the wild-type transporter (Fig. 2E, F, n = 7, ** p = 0.0019), suggesting the mutant has a reduced interaction with the adaptor and possibly some difficulty in ER output. Y707C immunocomplexes, in contrast, did not contain significantly different amount of Sec24D-Myc as compared to the wild-type (Fig. 2E, F, n = 7, ^ns^p = 0.1858).

### 3.3. Calnexin overexpression rescues the function and membrane arrival of the two transporter variants

We have previously reported that CNX overexpression facilitates the biogenesis and activity of wild-type GlyT2 (Arribas-Gonzalez et al. 2013). We expressed the transporters in COS7 cells together with increasing amounts of CNX and confirmed wild-type GlyT2 displays a progressive enhancement of expression and function (Fig. 3A, B). Since not all the GlyT2 mutants can be rescued by increasing CNX expression (Arribas-Gonzalez et al. 2015), we tested whether the mutants were amenable to rescue. Interestingly, the mutants showed even higher increases than the wild-type so that at the highest GlyT2 / CNX ratio (1:10), the two variants reached wild-type levels of glycine transport: 78.5 ± 3.3% and 94.9 ± 5.9% of the wild-type for A277T and Y707C, respectively, (n = 8, *** p [A277T 1: CNX 10] = 0.0003; *** p [Y707C 1: CNX 10] = 0.0007). This indicates the mutant phenotypes can be rescued.

**Figure 3.**
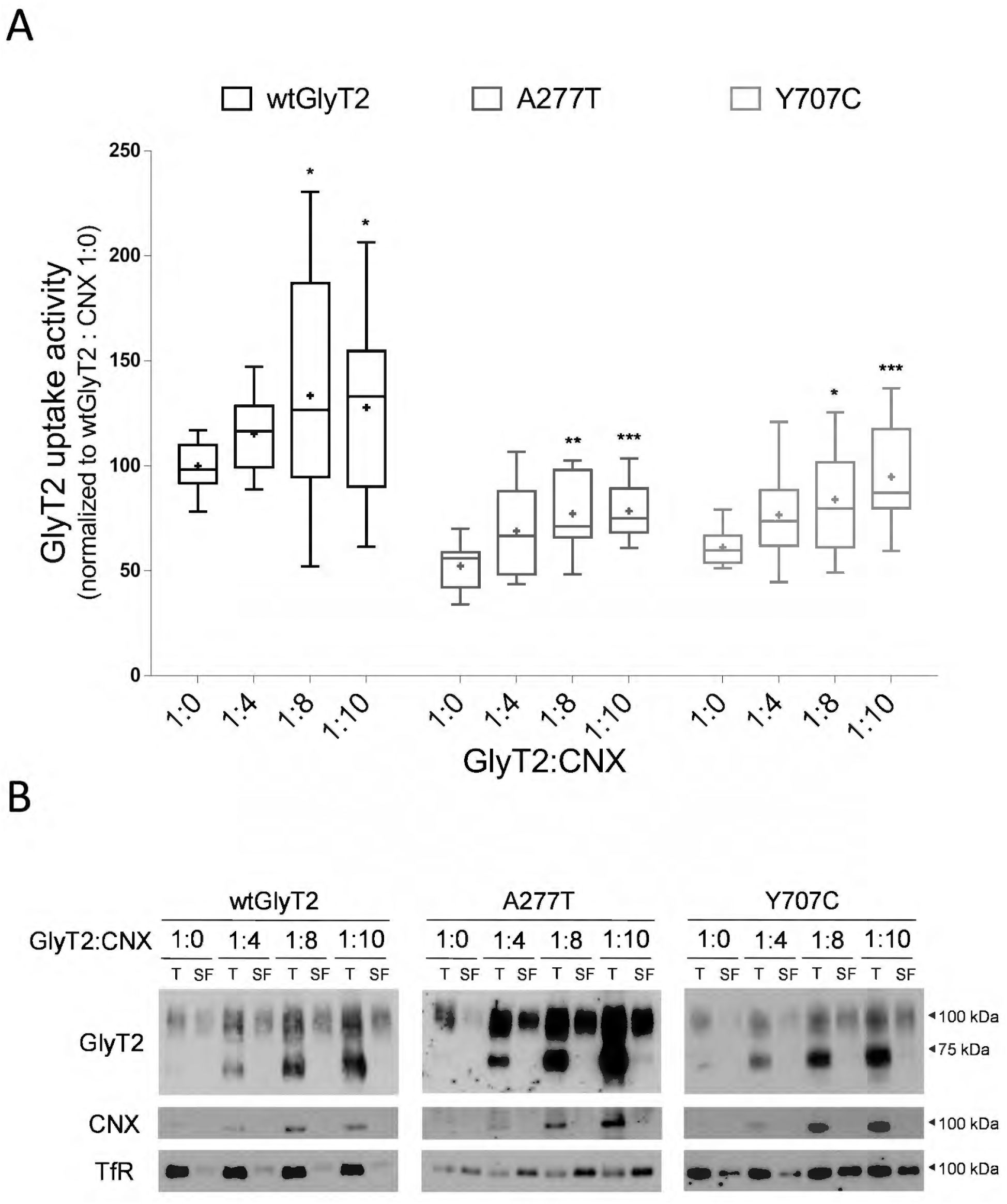
CNX overexpression rescues A277T and Y707C phenotype. COS7 cells transiently expressing CNX and either wtGlyT2, A277T or Y707C at the indicated ratios were subjected to [^3^H]-glycine transport assays (A) and surface biotinylation (B). **A)** Glycine transport rates of wtGlyT2 and mutants, normalized against wtGlyT2 in the absence of CNX. The statistical significance of the effect of CNX on transport was obtained by comparing glycine transport rates in the absence and in the presence of CNX for each variant. *p (wtGlyT2 1: CNX 8) = 0.0484, *p (wtGlyT2 1: CNX 10) = 0.0456, **p (A277T 1: CNX 8) = 0.0010, ***p (A277T 1: CNX 10) = 0.0003, *p (Y707C 1: CNX 8) = 0.0361, ***p (Y707C 1: CNX 10) = 0.0007; using Dunn’s multiple comparison test. n = 8. **B)** Representative western blot showing the effect of increasing amounts of CNX on surface biotinylated (SF) and total (T) fractions of wtGlyT2 and mutants. Transferrin receptor (TfR) immunoreactivity was used as loading control.

### 3.4. NAG increases the surface levels of wild-type GlyT2 in COS7 cells in a similar way as PBA

The facilitating action of CNX on the expression of GlyT2 variants associated with hyperekplexia stimulated the search for compounds that could rescue the phenotype of the mutants. We assayed four compounds as possible chemical chaperones: bupropion, ibogaine, ALX1393 and NAG. Bupropion and ibogaine are two atypical inhibitors of the serotonin and the dopamine transporters (SERT and DAT) that bind to the outward-facing conformation and whose efficacy as pharmacochaperones has been demonstrated in rescuing misfolded variants of these transporters (El-Kasaby et al. 2010; Beerepoot, Lam, and Salahpour 2016). ALX1393 is a reversible inhibitor of GlyT2 that binds to its open-out conformation (Vandenberg et al. 2016). NAG has been reported to be a non-competitive and reversible inhibitor of GlyT2 (Wiles et al. 2006). In addition, we used PBA as a positive control since its ability to improve the expression and function of wild-type GlyT2 was already demonstrated (Arribas-Gonzalez et al. 2015). In all cases, we treated COS7 cells expressing the transporters with the desired compound soon after the transfection, maintained the treatment during transporter synthesis, and removed it just before the assays (Fig. 4). In these conditions, as expected, 1 mM PBA increased surface GlyT2 around 40% as compared to vehicle (143.1 ± 14.4%; Fig. 4A, B, n = 4, *p = 0.0286) and promoted a modest (14%) enhancement of the transport activity (113.7 ± 4.7%; Fig. 4C, n = 16, *p = 0.0246). Treatments with bupropion and ibogaine at 1 μM, contrary to what was obtained for misfolded SERT and DAT mutants (El-Kasaby et al. 2010; Beerepoot, Lam, and Salahpour 2016), reduced the amount of GlyT2 present in the membrane (Fig. 4A) to a values of 71.8 ± 7.5%, n = 6, **p = 0.0022 (bupropion) and 59.2 ± 6.8%, n = 6, **p= 0.0022 (ibogaine) with respect to the treatment with vehicle. However, the reduction of membrane GlyT2 was not accompanied by a decrease in the transport activity (Fig. 4C, n = 26, ^ns^p (Bup) = 0.3020; ^ns^p (Ibo) = 0.2386). Regarding ALX1393, the treatment with a concentration as low as 10 nM produced about 20% increase in the surface fraction (119.5 ± 6.1%, as compared to vehicle, Fig. 4A, B, n = 4, *p = 0.0286), and a minimal increase in transport activity (105.7 ± 1.93%, as compared to vehicle; Fig 4C, n = 23, *p = 0.0489). Nevertheless, the most interesting compound was NAG, which at 10 μM produced 75% increase of the surface transporter (175.5 ± 35.8%; Fig. 4A, B, n = 4, *p = 0.0286), accompanied by a small (10%) increase in transport activity (109.5 ± 2.2%; Fig. 4C, n = 21, **p = 0.0046). This is comparable to PBA treatment.

**Figure 4.**
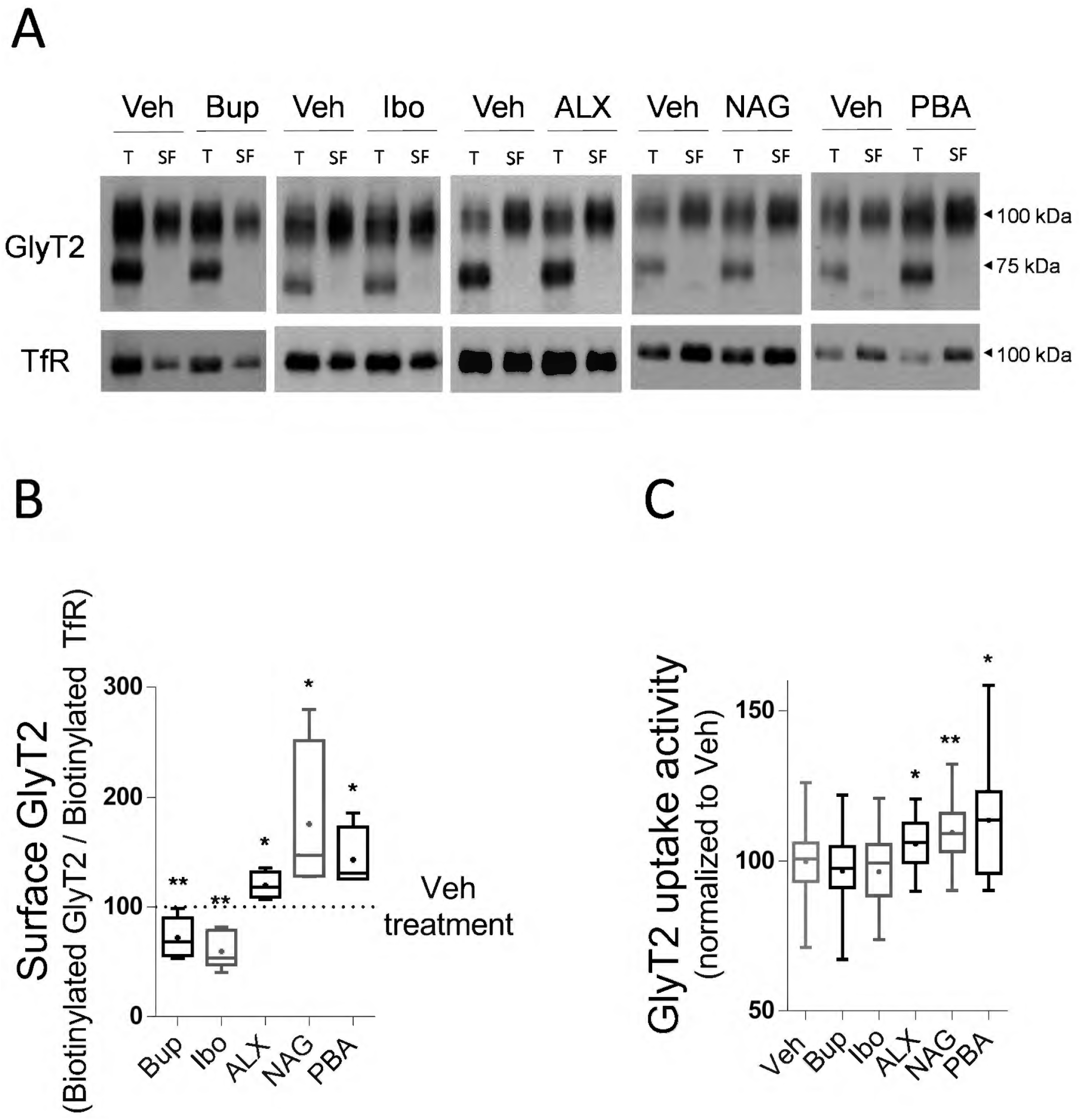
N-arachidonoyl glycine increases surface levels and transport activity of wtGlyT2 in COS7 cells. COS7 cells transiently expressing wtGlyT2 were treated with 1 μM bupropion (Bup), 1 μM ibogaine (Ibo), 1 nM ALX-1393 (ALX), 10 μM N-arachidonoyl glycine (NAG), 1 mM 4-phenylbutyric acid (PBA) or their vehicles (Veh) for 48 h. Then, cells were subjected to surface biotinylation (A, B) and [^3^H]-glycine transport assays (C). **A)** Representative western blot showing the effect of the assayed compounds on surface biotinylated (SF) and total (T) fractions of wtGlyT2. Transferrin receptor (TfR) immunoreactivity was used as loading control. **B)** Quantification of GlyT2 in the biotinlytated fraction using TfR as a loading control. The level of GlyT2 on the surface is expressed as a percentage of that obtained for the treatment with Veh in each case (indicated by the dashed line). **p (Bup) = 0.0022, **p (Ibo) = 0.0022, *p (ALX) = 0.0286, *p (NAG) = 0.0286, *p (PBA) = 0.0286; using Kolmogorov-Smirnov’s test. n (Bup) = 6, n (Ibo) = 6, n (ALX) = 4, n (NAG) = 4, n (PBA) = 4. **C)** Glycine transport rates of wtGlyT2. Gly transport is expressed as a percentage of that obtained for the treatment with Veh in each case. ^ns^p (Bup) = 0.3020, ^ns^p (Ibo) = 0.2386, *p (ALX) = 0.0489, **p (NAG) = 0.0046, *p (PBA) = 0.0246; using Kolmogorov-Smirnov’s test. n (Bup) = 26, n (Ibo) = 26, n (ALX) = 23, n (NAG) = 21, n (PBA) = 16.

### 3.5. NAG increases ER export of wild-type GlyT2 and decreases its interaction with CNX in COS7 cells

Despite the modest increase in activity, the large effect on the GlyT2 surface fraction observed after treatment with NAG gave indications of its potential to improve transporter expression. Therefore, we characterized the action of this compound. When applied from the extracellular medium, NAG inhibits the glycine transport by GlyT2 through direct interaction with the transporter expressed in *Xenopus laevis* oocytes (Wiles et al. 2006). We confirmed this inhibition takes also place in COS7 cells when the compound was added before or during the transport assay (Supplementary Fig. S3A, B), and not only during transporter biogenesis as above. NAG can bind to the transporter independently of glycine since NAG pre-treatment increases the inhibition potency. We discarded that residual NAG was responsible of the limited increase in glycine transport when the compound was delivered during transporter biogenesis as different washing procedures did not change the results (not shown). We also detected a small biphasic effect when NAG was added in the absence of glycine that produced some enhancement of the glycine transport at low µM concentrations (Supplementary Fig. S3B).

To further characterize the effects of NAG on wild-type GlyT2, we analyzed whether the treatment promoted changes in transporter export from the ER. COS7 cells expressing wild-type and treated for 48h with 10 μM NAG or vehicle were subjected to double immunocytofluorescence using antibodies against GlyT2 and the ER marker CNX (Fig. 5A). The colocalization degree was determined using Manders overlap coefficient. As a result, in NAG-treated cells, GlyT2 overlapped with CNX 21% less than in control cells (Fig. 5B, n = 31, *p = 0.0324), suggesting that somehow NAG promotes the release of GlyT2 from the ER. Next, to gain insights on the mechanism of action of NAG, we tested whether treatment with the compound during transporter biogenesis produced differences in the interaction of GlyT2 with CNX. As shown above, differences in the degree of CNX assistance could cause differences in membrane expression of the transporter. Therefore, we performed co-immunoprecipitation assays in COS7 cells after NAG treatment and found that the amount of GlyT2 co-immunoprecipitated with CNX was reduced to 57.2 ± 8.7% of the vehicle condition (Fig. 5C, D, n = 4, *p = 0.0286). Furthermore, it was observed that the interaction of GlyT2 with CNX was dependent on the NAG concentration used, obtaining the maximum reduction of the interaction with 10 µM (Supplementary Fig. S4). Interestingly, the treatment with 1 mM PBA also produced a decrease in the amount of GlyT2 co-immunoprecipitated with CNX, which reached 43.8 ± 19.9% (Fig. 5E, F, n = 3, *p = 0.0475), suggesting the two chaperones somehow restrict the interaction CNX-GlyT2 perhaps with related mechanisms.

**Figure 5.**
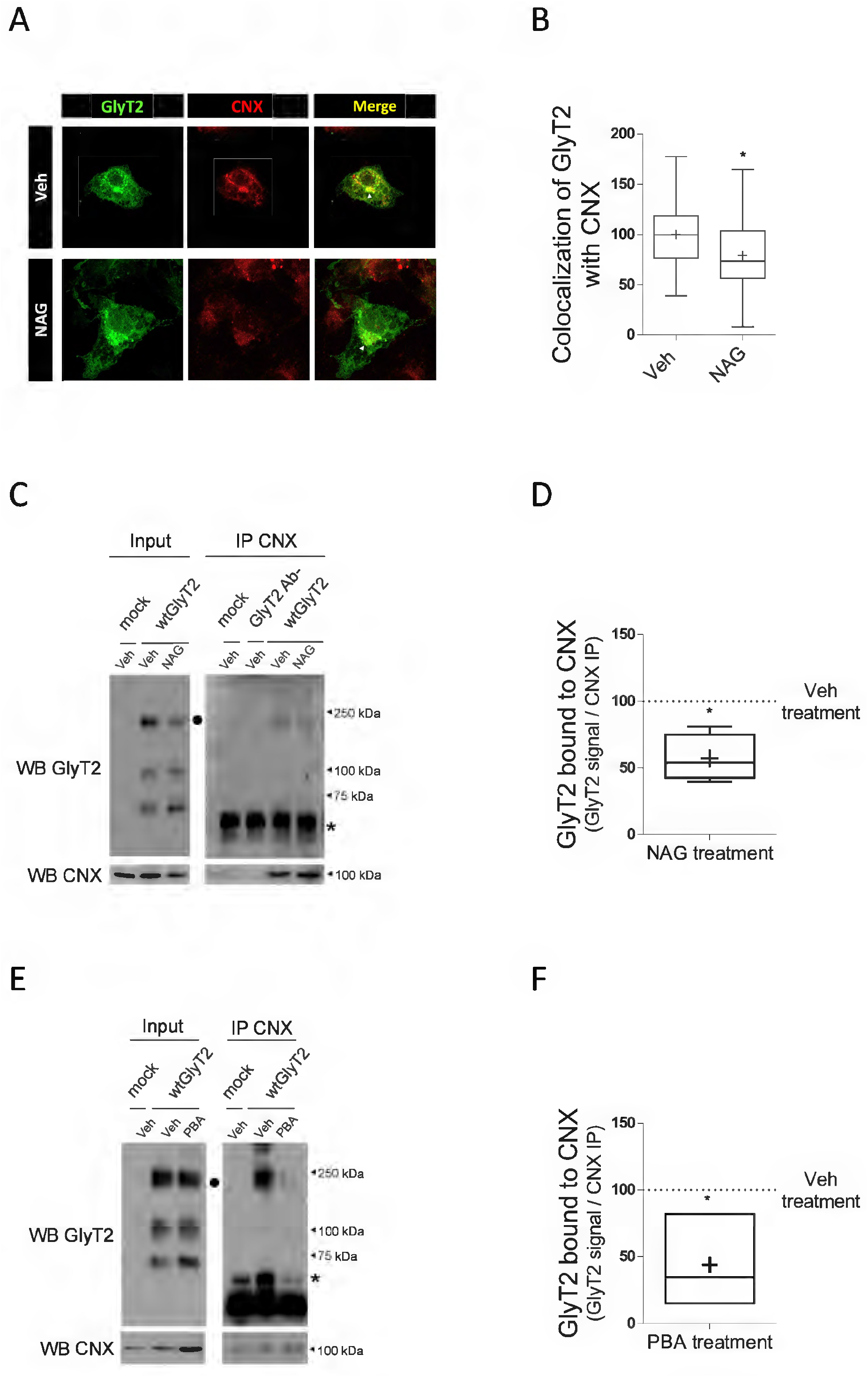
N-arachidonoyl glycine decreases the colocalization and interaction of GlyT2 with CNX in COS7 cells. **A)** COS7 transiently expressing wtGlyT2 were treated with 10 μM N-arachidonoyl glycine (NAG) or vehicle (Veh) for 48 h and immunolabeled for GlyT2 (green) and the endoplasmic reticulum marker CNX (red). Arrowheads (△) indicates areas showing most colocalization. **B)** Colocalization of wtGlyT2 with CNX was quantified using Mander’s overlap coefficient and normalized against Veh. *p = 0.0324; n (Veh) = 25, n (NAG) = 31. **C)** COS7 cells transiently transfected and treated as in (A) were subjected to CNX immunoprecipitation using anti-CNX made in rabbit and the immunocomplexes were analyzed by western blot to detect CNX and GlyT2 (using anti-GlyT2 made in rat). 250 kDa aggregates of inmature GlyT2 are indicated with a black circle (●); unspecific 50 kDa bands are indicated with an asterisk (*). **D)** Quantification of the amount of GlyT2 bound to CNX after treatment with 10 μM NAG expressed as a percentage of the amount determined after Veh treatment (indicated by the dashed line). *p = 0.0286, using Komogorov-Smirnov’s test. n = 4. **E)** COS7 cells transiently transfected as in (A) were treated with 4-phenylbutyric acid (PBA) 1 mM or its vehicle (Veh) for 48 h and subjected to immunoprecipitation against CNX. Then, the immunocomplexes were analyzed by western blot to detect GlyT2 and CNX. **F)** Quantification of the amount of GlyT2 bound to CNX after treatment with 1 mM PBA expressed as a percentage of the amount determined after Veh treatment (indicated by the dashed line). *p = 0.0475, using unpaired Student’s t test. n = 3. Mock: untransfected cell sample. GlyT2 Ab-: GlyT2 transfected but precipitation without antibody.

### 3.6. NAG rescues the activity and surface expression of the two hyperekplexia-associated variants in primary neurons

Since NAG was able to exert chaperone action on wild-type GlyT2, we determined if it had the ability to rescue the GlyT2 mutants. For these experiments we used a system closer to the endogenous GlyT2 and expressed the transporters in primary neurons of rat cerebral cortex where the two mutants behave as in COS7 cells, showing reduced transport activity and membrane expression (Fig. 6A-C). The glycine transport by A277T was 58.6 ± 4.0%, with respect to the wild-type (Fig. 6A, n = 32, ****p <0.0001), and its membrane expression was reduced to 40.1 ± 7.7% (Fig. 6C, n = 13, ** p = 0.0094). Y707C showed an activity and membrane expression of 54.2 ± 4.3% (Fig. 6B, C, n = 33, ****p <0.0001) and 31.4 ± 11.15 % (Fig. 6C, n = 7, *p = 0.0186), as compared to the wild-type.

**Figure 6.**
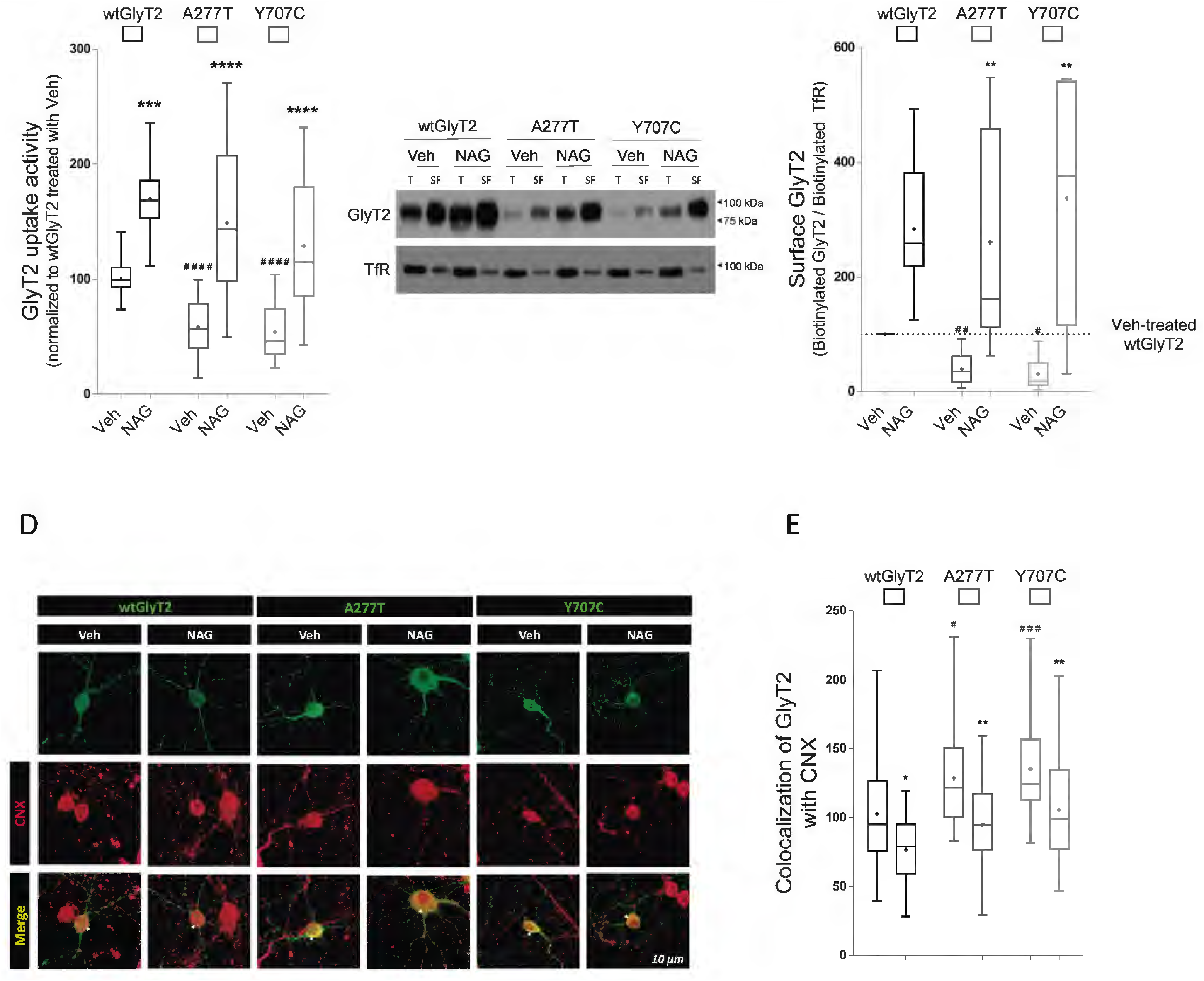
N-arachidonoyl glycine rescues A277T and Y707C phenotype in primary cortical neurons. Primary cortical neurons transiently expressing wtGlyT2 (wild-type), A277T or Y707C were treated with 10 μM N-arachidonoyl glycine (NAG) or vehicle (Veh) for 48 h. Then, neurons were subjected to [^3^H]-glycine transport (A), surface biotinylation assays (B, C) and double immunofluorescence against GlyT2 and CNX (D, E). **A)** Glycine transport rates as a percentage of wtGlyT2 treated with Veh. (#) statistically significant difference when comparing the Veh condition of the indicated mutant with Veh of wtGlyT2. ^# # # #^p < 0.0001, using Dunn’s multiple comparison test. n (wtGlyT2 + Veh) = 46, n (A277T + Veh) = 32, n (Y707C + Veh) = 33. (*) statistically significant difference when comparing the Veh and the NAG condition of each variant. ****p (wtGlyT2 + NAG) = 0.0004, ****p (A277T + NAG) < 0.0001, ****p (Y707C + NAG) < 0.0001, using Dunn’s multiple comparison test. n (wtGlyT2 + NAG) = 25, n (A277T + NAG) = 18, n (Y707C + NAG). **B)** Representative western blot showing the effect of NAG on surface biotinylated (SF) and total (T) fractions of GlyT2. Transferrin receptor (TfR) immunoreactivity was used as loading control. **C)** Quantification of GlyT2 biotinlytated fraction using TfR as loading control normalized to the corrected signal of GlyT2 treated with Veh (indicated by the dashed line). (#) significantly different from wtGlyT2 treated with Veh. ^# #^p (A277T + Veh) = 0.0094, ^#^p (Y707C + Veh) = 0.0186, using Dunn’s multiple comparison test. (wtGlyT2 + Veh) = 15, n (A277T + Veh) = 13, n (Y707C + Veh) = 7. (*) significantly different when comparing the NAG and the Veh condition of each variant. **p (A277T + NAG) = 0.0033, **p (Y707C + NAG), using Dunn’s multiple comparison. n (wtGlyT2 + NAG) = 8, n (A277T + NAG) = 5, n (Y707C + NAG) = 5. **D)** Representative confocal images showing the effect of NAG on the GlyT2 overlap with ER marker CNX. Arrowheads (△) indicates areas showing most colocalization. **E)** Colocalization with CNX, quantified using Mander’s overlap coefficient, are normalized against wtGlyT2 treated with Veh. (#) colocalization with CNX significantly different from that of wtGlyT2 treated with Veh. ^#^p (A277T) = 0.0234, ^# #^p (Y707C) = 0.0025; using Holm-Sidak’s multiple comparison test. n (wtGlyT2 + Veh) = 32, n (A277T + Veh) = 29, n (Y707C + Veh) = 29. (*) colocalization with CNX significantly different when comparing the NAG and the Veh condition of each variant. **p (A277T + NAG) = 0.0016, **p (Y707C + NAG) = 0.0075); using Holm-Sidak’s multiple comparison test. n (wtGlyT2 + NAG) = 31, n (A277T + NAG) = 33, n (Y707C + NAG) = 32.

Neurons (between DIV 5 and 7) that had been transfected to express every transporter mutant were incubated with 10 μM NAG added immediately after transfection and maintained the subsequent 48h. Significant increases were observed in the activity and surface expression of the mutants (Fig.6A, B). Glycine transport and surface expression of A277T increased in the presence of NAG to 148.8 ± 14.8% (Fig. 6A, n = 18, ****p <0.0001) and 260.5 ± 87.2% (Fig. 6C, n = 5, **p = 0.0033) from the wild-type, respectively; while, in the case of Y707C, the transport activity was 129.3 ± 13.8% (Fig. 6A, n = 18, ****p <0.0001) and the membrane expression was 337.4 ± 99.0% (Fig. 6C, n = 5, **p = 0.0028), with respect to the wild-type.

Finally, we studied whether NAG treatment influenced the degree of ER retention of the GlyT2 mutants. Rat brain primary neurons from cerebral cortex were transfected with the cDNAs of the transporters and treated with 10 μM NAG during the 48h post-transfection. After treatment, the neurons were subjected to double immunocytofluorescence using antibodies against GlyT2 and CNX. The degree of retention in the ER was determined using the Manders colocalization coefficient. As expected, in the untreated neurons the mutants were retained in the ER, showing 30% higher colocalization with CNX than the wild-type transporter (Fig. 6D, E): A277T and Y707C colocalized with CNX 128.3 ± 6.2% and 135.2 ± 6.9%, respectively (Fig. 6E, n (A277T + Veh) = 29, n (Y707C + Veh) = 29, *p (A277T) = 0.0234, **p (Y707C) = 0.0025), values very close to those determined in COS7 cells (Fig. 1H). In the neurons, NAG reduced the degree of overlap with CNX of the mutants to the levels of the wild-type transporter under control conditions. CNX overlap of A277T and Y707C reached 94.9 ± 5.0% and 105.6 ± 7.0% of the overlap shown by the wild-type transporter, respectively, Fig. 6D, E, n (A277T + NAG) = 33, n (Y707C + NAG) = 32, **p (A277T) = 0.0016, **p (Y707C) = 0.0075. In conclusion, these results suggest that NAG increases the activity of the A277T and Y707C mutants by reducing ER retention and subsequently increasing plasma membrane expression.

## 4. Discussion

In this work we have studied the intracellular trafficking of two GlyT2 variants previously associated to hyperekplexia that contain missense mutations affecting several functions of the transporters (Carta et al. 2012; Gimenez et al. 2012). Here we provide new information demonstrating the two variants show ER retention and more intense association with the ER chaperone CNX than the wild-type transporter. One of the two mutants (A277T) have also a defective association with the ER export adaptor Sec24D, probably due to folding alterations. This substitution lays close to the Na3 site of the transporter, a region with robust allosteric properties and conceivably important for the folding (Benito-Munoz et al. 2018). However, in the other variant (Y707C), the substitution, located in the C-t portion of TM11 at the extracellular face of the transporter (Gimenez et al. 2012), may have a minor role in the folding of the protein core. In agreement with this idea, we proved A277T but not Y707C displays higher level of ubiquitination, suggesting the mutant is more prone to suffer ERAD. Therefore, this quality control may eliminate higher levels of this mutant than of Y707C or wild-type. Even so, globally, the presence of the two mutants in the plasma membrane is similarly reduced, so that the Y707C mutant must undergo additional retention in other components of the endomembrane system.

Here we have also proven the phenotype of the two mutants can be rescued by CNX overexpression and by using chemical chaperones. We tested several compounds with potential to improve the function of GlyT2. Among them, the atypical monoamine inhibitors bupropion and ibogaine promoted opposite effects to those obtained by other authors for wild-type DAT (Beerepoot, Lam, and Salahpour 2016). These compounds inhibit monoamine transporters by blocking the proteins in inward-open conformation and facilitate the maturation acting as pharmacochaperones. This action has led to the proposal that, during folding, SLC6 transporters have to adopt the inwardly-directed conformation for the acquisition of the native folding (Freissmuth, Stockner, and Sucic 2018; Bhat, Newman, and Freissmuth 2019; Asjad et al. 2017). For GlyT2, instead, these compounds decreased surface levels although the GlyT2 fraction in the membrane of bupropion or ibogaine-treated cells, despite being inferior, had the same ability to transport glycine as the membrane fraction of untreated cells. Although the affinity of these compounds for GlyT2 is unknown, these data would support the hypothesis that acquiring an inward-open conformation facilitates folding. However, since treatment with these compounds did not improve the global transport capacity of the cells expressing GlyT2, they are unpromising compounds at least in its actual form.

The GlyT2 transport inhibitor, ALX1393, produced significant increase in the amount of plasma membrane GlyT2 but minimal increase in its activity. Although this compound binds and inhibits GlyT2 with nM affinity (Xu, Gong, and Xu 2005; Mingorance-Le Meur et al. 2013), the possibility that it acts on GlyT2 folding in the ER is, however, scarce. First, it is not known if ALX1393 can cross the plasma membrane of cells, but it seems not very likely taking into account its reduced capability of blood-brain barrier permeation (Mingorance-Le Meur et al. 2013). Secondly, even if it can reach the ER lumen, it would not be able to interact with GlyT2 in that context, as ALX1393 binding to the outward-open conformation requires sufficient Na^+^ concentration to form the Na1 site (Benito-Munoz et al. 2018), and the concentration of the cation in the ER lumen is minimal (Bhat, Newman, and Freissmuth 2019). Perhaps this is the reason why the so-called typical DAT inhibitors, which have an inhibition mechanism similar to that of ALX1393, were not effective in promoting folding (Beerepoot, Lam, and Salahpour 2016). The increase in plasma membrane GlyT2 after treatment with ALX1393, therefore, may be the result of the binding of the inhibitor to the cell surface transporter, as the extracellular Na^+^ concentration allows the interaction. The study of GlyT2 exocytosis in the presence of ALX1393 may help to understand the mechanism behind this phenomenon.

Out of all the compounds tested in this work, the most promising was NAG. The treatment of GlyT2 mutants expressed in rat cortical neurons with NAG reduced transporter binding to CNX and ER retention, and therefore, greatly increased transport activity and membrane expression. Decreased interaction of cargo proteins with CNX or with other molecular chaperones was reported by several authors working with pharmacochaperones. Ibogaine diminished the interaction of CNX with a SERT mutant (El-Kasaby et al. 2010) and noribogaine treatment reduced the interaction of HSP70 with a group of DAT mutants associated with childhood and juvenile parkinsonism (Asjad et al. 2017). A decrease of the chaperone-transporter interaction can be mechanistically achieved by facilitating folding through the specific binding of the pharmacochaperone to the transporter, what subsequently reduced the number of chaperone-cargo complexes. As NAG is likely to reach the ER lumen due to its structure (Burstein 2018), the inhibition of CNX action could be due to the direct binding of the compound to GlyT2, an interaction that may somehow block or compete with CNX assistance. Transport inhibitors trapping the transporters in a certain conformation, may exert its action by this mechanism of pharmacochaperoning (Bhat, Newman, and Freissmuth 2019; Asjad et al. 2017). Our data do not exclude NAG acts directly on the transporter conformation, although the fact that the optimal NAG concentration for increasing GlyT2 expression and function (10 μM) was much lower than the IC_50_ for GlyT2 inhibition (67 ± 23 μM), conflicts this possibility. A related possibility is that NAG improves the folding of GlyT2 through a less specific interaction, that is, acting as a chemical chaperone. In this sense, GlyT2 treatment with NAG gave similar results in terms of activity, membrane expression and interaction with CNX than the treatment with PBA, a well-established chemical chaperone (Arribas-Gonzalez et al. 2015; Kaur et al. 2018). The two compounds may act in a similar way. NAG, as well as PBA, contain hydrophobic regions that may create a hydrophobic context around GlyT2 that facilitates folding. The clear biphasic effect in the response of GlyT2 to externally applied NAG with a slight stimulation at concentrations around 10 μM, may suggest a stabilizing action and perhaps a chaperone role.

Without ruling out the previous option, it should not be neglected that NAG does not have to improve GlyT2 folding, since the lower interaction of GlyT2 with CNX may be due to a blockage of chaperone assistance (Egan et al. 2002). In this case, treatment with NAG would favor the arrival of GlyT2 to the plasma membrane by promoting the transporter to escape the quality control exerted by CNX. The loosening of CNX-GlyT2 interaction could be also due to some indirect mechanism. For example, CNX uses Ca^2+^ as a cofactor for protein binding (Hebert and Molinari 2007; Ellgaard and Frickel 2003). The depletion of the ER intraluminal Ca^2+^ levels can impair the interaction, increasing surface expression of the CNX client. This was proposed for the CFTR ΔF508 mutant after inhibiting ER Ca^2+^ ATPase (SERCA) with thapsigargin (Egan et al. 2002). In this sense, it is worthy to mention that NAG may trigger several signaling actions, some of which can affect Ca^2+^ concentrations within the ER and that it would be interesting to explore in the near future (Kohno et al. 2006; Deak et al. 2013). Anyhow, in this study we have shown that the presence of NAG during transporter biogenesis greatly improves the ER output, membrane expression and activity of GlyT2, what makes NAG a compound with real capacity to rescue in vitro the phenotype of GlyT2 mutants associated with hyperekplexia that have retention problems in ER. Maturation deficits are not only known to cause hyperekplexia in *SLC6A5* patients, but also in mutations affecting the glycine receptor (GlyRα1). Whether NAG could have a role in vivo deserves further research.

## 5. Conclusion

The present results show that two GlyT2 hyperekplexia missense mutations with defective traffic are partially retained in the ER and their interaction with CNX is increased. Variant A277T has export difficulties and increased ubiquitination levels what suggest it is a preferential subject of ER-associated degradation. The membrane arrival and function of the two mutant transporters can be increased by CNX overexpression and chemical chaperones indicating their phenotype is amenable to rescue. Among several compounds tested, we found that NAG, an arachidonic acid derivative with inhibitory action on GlyT2, can rescue the trafficking defects of the two variants in heterologous cells and rat brain cortical neurons. NAG treatment during transporter biogenesis loosens the quality control exerted by CNX increasing surface expression and therefore transport activity. N-arachidonoyl glycine and its derivatives have potential for hyperekplexia therapy.

## Abbreviations

ALX: ALX1393
Bup: bupropion
CBMSO: Centro de Biología Molecular Severo Ochoa
CEI-UAM: Comité de Ética de la Investigación UAM
CFTR: cystic fibrosis transmembrane conductance regulator
CNX: calnexin
COPII: coatomer protein II
COS: CV-1 (simian) in origin and carrying the SV40 genetic material
DAT: dopamine transporter
DMEM: Dulbecco’s modified Eagle’s medium
DTT: dithiothreitol
E-cadh: E-cadherin
ECL: enhanced chemiluminescence
EDTA: ethylenediaminetetraacetic acid
Endo H: endoglycosidase H
ER: endoplasmic reticulum
ERAD: endoplasmic reticulum-associated degradation
FBS: fetal bovine serum
GABA: γ-aminobutyric acid
GlyR: glycine receptor
GlyT2: glycine transporter 2
HSP70: heat-shock protein 70
Ibo, ibogaine: 
IgG: Immunoglobulin G
MDCK: Madin-Darby canine kidney
MEM: minimum essential medium
mock: non-transfected cells
NAG: N-arachidonoyl glycine
NFPS: N-[3-([1,1-Biphenyl]-4-yloxy)-3-(4-fluorophenyl)propyl]-N-methylglycine
OMIM: Online Mendelian Inheritance in Man
NHS: N-Hydroxysuccinimide
PBA: 4-phenylbutyric acid
PBS: phosphate buffered saline
PMSF: phenylmethylsulfonyl fluoride
PNGase F: peptide: N-glycosidase F
PGA: protein G agarose
RIPA: radioimmunoprecipitation assay
RT: room temperature
SDS-PAGE: sodium dodecyl sulfate polyacrylamide gel electrophoresis
SERCA: ER Ca2+ ATPase
Serotonin transporter: 
SERTSERT: serotonin transporter
SLC: solute carrier
TfR: transferrin receptor
TM: transmembrane domain
Tub: tubulin
Ub: ubiquitin
WB: western blot.

## Author contributions

Conceived the work: B.L.-C. Performed the experiments: A.R-M. and E. M. Analyzed the data: B.L.-C., A.R-M. and C.A. Wrote the paper: B.L.-C. and A.R-M.

## Acknowledgments

The authors are grateful to FJ Díez-Guerra and JM Cuezva for generous support. The confocal microscopy facility at the CBMSO (Madrid, Spain) is also acknowledged for valuable and expert help with the confocal microscopy work. Enrique Núñez is acknowledged for excellent technical assistance.

