## Supplementary Fig. S1 for "Rescue of two trafficking-defective variants of the neuronal glycine transporter GlyT2 associated to hyperekplexia"

Andrés de la Rocha-Muñoz[a,b], Elena Melgarejo[a,b], Enrique Núñez[a,b,c], Carmen Aragón[a,b,c] and Beatriz López-Corcuera[a,b,c*]

[a]Departamento de Biología Molecular Universidad Autónoma de Madrid. Spain.
[b]Centro de Biología Molecular "Severo Ochoa" Consejo Superior de Investigaciones Científicas-Universidad Autónoma de Madrid. Spain.
[c]IdiPAZ-Hospital Universitario La Paz, Universidad Autónoma de Madrid, Spain.

### Supplementary Information

### Supplementary Figures

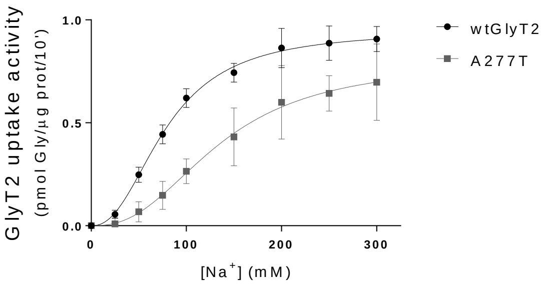


**Supplementary Figure S1. A277T presents altered $Na^{+}$ dependence of glycine transport.** COS7 cells transiently transfected with wtGlyT2 or A277T were subjected to [$^{3}$H]-glycine transport assays at increasing concentrations of extracellular NaCl (isotonically replaced by choline chloride).

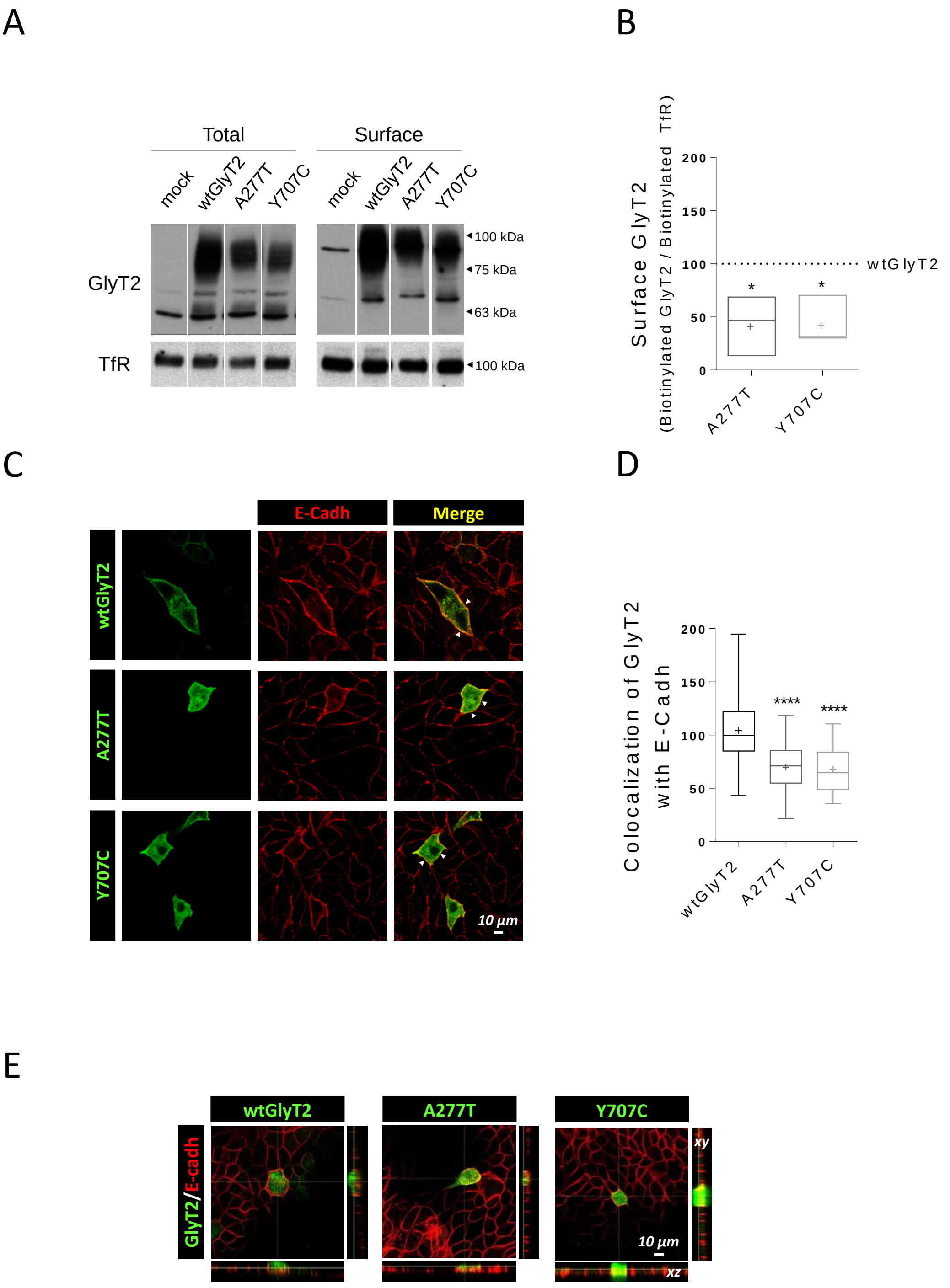


**Supplementary Figure S2. A277T and Y707C also present defective trafficking properties in MDCK II cells. A)** MDCK II cells transiently expressing wtGlyT2 (wild-type), A277T or Y707C were subjected to surface biotinylation: surface biotinylated (SF) and total (T) transporter fractions from the lysate were analyzed by western blot. Transferrin receptor (TfR) immunoreactivity was used as loading control. **B)** Quantification of transporter biotinlytated fraction using TfR as loading control (percentage to the corrected signal of wtGlyT2, indicated by the dashed line). *p (A277T) = 0,0268, *p (Y707C) = 0,0288, using Dunnet's multiple comparison test. n = 3. **C)** MDCK II cells transiently transfected as in (A) were immunolabeled for GlyT2 (green) and the plasma membrane marker E-cadherin (E-cadh) (red). Arrowheads (△) indicates areas showing most colocalization. **D)** Colocalization of transporters with E-cadh was quantified using Mander's overlap coefficient, which determines the degree of colocalization between two channels: the proportion of transporter (green) that colocalizes with E-cadh (red). Colocalization levels are normalized against those shown by wtGlyT2. ****p < 0.0001, using Dunn's multiple comparison test. n (wtGlyT2) = 47, n (A277T) = 35, n (Y707C) = 32. **E)** MDCK II cells transiently transfected as in (A) were plated on cell culture filter inserts and grown to confluence. Then, samples were inmulolabeled as in (C) and examined by laser scanning confocal microscopy. *Central panel*, en face views. *Right panel*, xy cross-sections. *Botton panel*, xz cross-sections

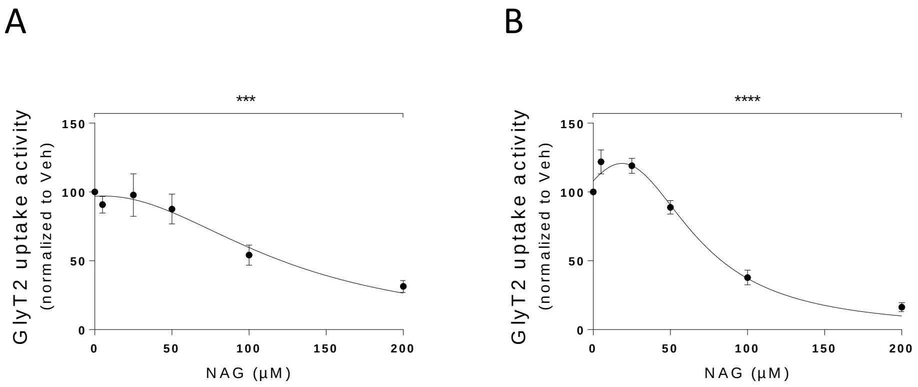


**Supplementary Figure S3. The presence of N-arachidonoyl glycine during [$^3$H]-glycine transport assays in COS7 cells inhibits glycine transport rates of wtGlyT2.** COS7 cells transiently expressing wtGlyT2 were subjected to [$^3$H]-glycine transport assays in the presence of increasing concentrations of N-arachidonoyl glycine (NAG) (0-200 µM). **A)** Glycine transport rates of wtGlyT2 in the presence of NAG without a previous incubation with the drug. ***p = 0.0009, using Kruskall-Wallis multiple comparison test. n = 6. **B)** Glycine transport rates of wtGlyT2 in the presence of NAG after a 10 min pre-incubation with the drug at the same concentrations used during the assay. ****p = 0.0001, using Kruskall-Wallis multiple comparison test. n = 15.

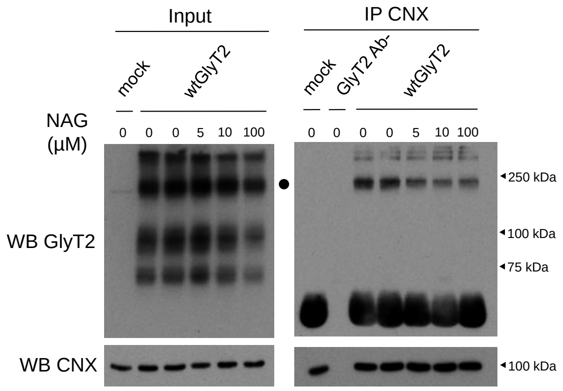


**Supplementary Figure S4. N-arachidonoyl glycine decreases the interaction of GlyT2 with CNX in a dose-dependent manner.** COS7 cells transiently transfected with wtGlyT2 were treated with N-arachidonoyl glycine (NAG) at the indicated concentrations or its vehicle (Veh) for 48 h and subjected to CNX immunoprecipitation. Then, the immunocomplexes were analyzed by western blot to detect GlyT2 and CNX.
